# QuickStitch for seamless stitching of confocal mosaics through high-pass filtering and recursive normalization

**DOI:** 10.1101/075440

**Authors:** Pavel A. Brodskiy, Paulina M. Eberts, Cody Narciso, Jochen Kursawe, Alexander Fletcher, Jeremiah J. Zartman

**Affiliations:** Department of Chemical and Biomolecular Engineering, University of Notre Dame, 182 Fitzpatrick Hall, Notre Dame, IN 46556, USA.; Mathematical Institute, University of Oxford, Andrew Wiles Building, Radcliffe Observatory Quarter, Woodstock Road, Oxford, OX2 6GG, UK.; School of Mathematics and Statistics, University of Sheffield, Hicks Building, Hounsfield Road, Sheffield, S3 7RH, UK.; Bateson Centre, University of Sheffield, Sheffield, S10 2TN, UK

**Author notes:** These authors contributed equally to this work (Pavel A. Brodskiy, Paulina M. Eberts).

**Keywords:** confocal microscopy, image processing, flat-field correction, uneven illumination, epithelial segmentation, open-source

## Abstract

Fluorescence micrographs naturally exhibit darkening around their edges (vignetting), which makes seamless stitching challenging. If vignetting is not corrected for, a stitched image will have visible seams where the individual images (tiles) overlap, introducing a systematic error into any quantitative analysis of the image. Although multiple vignetting correction methods exist, there remains no open-source tool that robustly handles large 2D immunofluorescence-based mosaic images. Here, we develop and validate QuickStitch, a tool that applies a recursive normalization algorithm to stitch large-scale immunofluorescence-based mosaics without incurring vignetting seams. We demonstrate how the tool works successfully for tissues of differing size, morphology, and fluorescence intensity. QuickStitch requires no specific information about the imaging system. It is provided as an open-source tool that is both user friendly and extensible, allowing straightforward incorporation into existing image processing pipelines. This enables studies that require accurate segmentation and analysis of high-resolution datasets when parameters of interest include both cellular-level phenomena and larger tissue-level regions of interest.

## INTRODUCTION

In immunofluorescence microscopy studies of tissue-scale morphology, mosaics comprised of multiple fields of view are often required to obtain high-resolution data necessary to study cell-level properties. For example, quantitatively connecting cell shape parameters to tissue-level morphogenetic processes is a key challenge in developmental biology (1–3). Due to the non-uniform illumination present in all optical equipment, objects located near the edges of fluorescence micrographs have a lower signal intensity than those in the center of the field of view (4). This causes stitched montages to have visible seams (vignetting) that reduce the accuracy of image data (5). While commercial systems are increasingly reducing the effect of vignetting by providing more uniform illumination, vignetting is never entirely eliminated. A solution is needed to generate seamless mosaics of fluorescent images of large tissues with sparse background, such as histological samples.

To address this, we have developed QuickStitch, an open-source, freely available, computationally efficient image processing tool that applies a high-pass filter either through a Fast Hartley Transform (FHT) or by Gaussian filtering to individual tiles, then fits normalization parameters to minimize the sum of squared error (SSR) in the overlap. The Fast Fourier Transform (FFT) is a computational algorithm commonly used to decompose sequences into a sum of sinusoidal functions (6). FFT high-pass filtering can be approximated by using an FHT band-pass filter to remove low-frequency objects affecting the entire field of view, such as vignetting artifacts, while preserving the high-frequency features that comprise the raw image. To demonstrate the tool’s efficacy, we validated QuickStitch on whole-mount *Drosophila* embryos.

## BACKGROUND

Several flat-field correction methods are commonly used to approximate the original pattern of a signal from the raw data collected. The flat-field correction equation for vignetting is given by

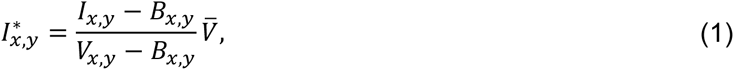

where *I* and *I\** are the original and corrected image matrices respectively; *V* is the vignetting mask, which is a matrix of the same size as the image matrix; *⊽* is its spatial average; and *B* is the background intensity (7). The indices *x* and *y* in eq. (1) represent the position of pixels in the *x* and *y* directions. In practice, *B* is frequently assumed to be constant across the field of view, hence does not contribute to vignetting artifacts.

A variety of methods used for approximating *V* and *B* are summarized below:

### Fluorescent correction slide

A common recourse for flat-field correction is to obtain an image of a blank area on a slide to utilize as a vignetting mask (7–10). This image is assumed to be the inverse of *V,* while *B* is taken to be the zero matrix. This approach is applicable for brightfield microscopy; however, in the case of fluorescence microscopy, a fluorescent slide must be used because the empty region of the glass slide does not yield a fluorescence signal (7). Furthermore, neglecting *B* causes artifacts in the interpretation of fluorescence data, which results in the over or under-estimation of regional concentrations (11). To avoid this, *B* may instead be an image taken while the lasers are inactive. Another weakness of this method is the assumption that the vignetting mask is the same for all images, regardless of the signal intensity (7–9).

### Background segmentation

Background segmentation techniques can be used to correct for vignetting after image acquisition via the creation of a vignetting mask from background information, typically by averaging or taking the median of background pixels (12,13). However, this information may not be present if the sample takes up the whole viewing area, or may be difficult to segment in whole-tissue mosaics given that different tissue types and cell cultures frequently require tailored segmentation methods. Such background detection methods assume that the vignetting mask is dependent only on position and not intensity, enabling the user to obtain a background image from a composite of multiple fields of view (12,14). As a result, this requires segmentation to separate the background from the foreground, which is difficult to do for features of varying size, because an entirely new segmentation algorithm is needed for each application. This method also requires many images to approximate the vignetting parameters. This method attempts to construct a background slide image from regions of the image that do not contain objects, meaning that while it can be used to approximate *B,* approximation of *V* is challenging. Open-source implementations of this method are available, but are of limited applicability to images of large tissues that expose only small sections of background (14,15).

### Physical (parametric) principles

Another class of methods for vignetting correction is to solve for the vignetting mask using analytical solutions derived from the physics governing vignetting (16,17). Generally, focal length, principal point, aspect ratio, and skew of the lens must either be provided or measured from reference material. Good approximations of the vignetting function can even be made without a reference slide if the geometry of the optical instrumentation is known such that these parameters can be obtained. Since precise specifications that govern every piece of optical equipment involved in the imaging are needed, this method is most feasible for applications by the companies that supply the optical equipment. Highly informed parametric methods are also often slower than other methods. Further, it is impossible for a parametric method to account for all sources of vignetting (16,17). This method is not a practical solution for end-users or for custom equipment, but is often implemented by equipment manufacturers.

### Image averaging methods

For image averaging methods, an “average image” is generated from a set of images (18). The inverse of the average is taken to be the vignetting mask. This method does not require a reference slide; however, it assumes that objects in the image are uniformly distributed, meaning that a large number of reference images are needed if the landmarks are not sufficiently homogeneous (18).

### High-pass filtering methods

Frequency-filtering methods decompose an image into a sum of images of various spatial frequencies. Low frequency features such as the background are separated from high frequency effects such as details and noise. Although FFT is more frequently employed in image de-noising algorithms (19), here we utilize FFT filtering of the tiles to remove features lower in frequency than the cutoff frequency, which is greater than the frequency of vignetting across an empty field, but less than the frequency of the largest object. Consequently, a major limitation of this approach is that vignetting effect must have a larger period than other features in the images; however, this approach does not suffer from the limitations imposed by other approaches which require many images, segmentation, or reference images. This results in the removal of vignetting artifacts because vignetting is a low-frequency phenomenon.

### Need for a normalization method

In the course of obtaining high-resolution, multi-tiled immunofluorescent image data sets of late stage *Drosophila* embryos, we found that visible seams caused by vignetting were present. We tested several vignetting correction methods previously described in the literature in an effort to correct for vignetting to produce clearly defined mosaics with image features of variable characteristic sizes. We concluded that none of the approaches satisfactorily correct for vignetting. In particular, we compared our proposed method to the gold standard rhodamine fluorescence slide method, and found that our method was better able to qualitatively eliminate the seams arising from vignetting.

## MATERIALS AND METHODS

### Immunohistochemistry

A *Drosophila* line expressing GAL4 under the engrailed *(en)* promotor and CD8::GFP under the UAS promotor was used in preparing Figure 2. The immunohistochemistry (IHC) protocol was based on previously described experiments (20,21) optimized for dpERK labeling with rabbit anti-dpERK (1:100, Cell Signaling), rat DCAD2 (1:100, DSHB) and DAPI (5 μg/ml, Invitrogen DU1306), with goat anti-rat IgG 561 (1:500, Invitrogen), and goat anti-rabbit IgG 647 (1:500, Invitrogen).

**Figure 2:**
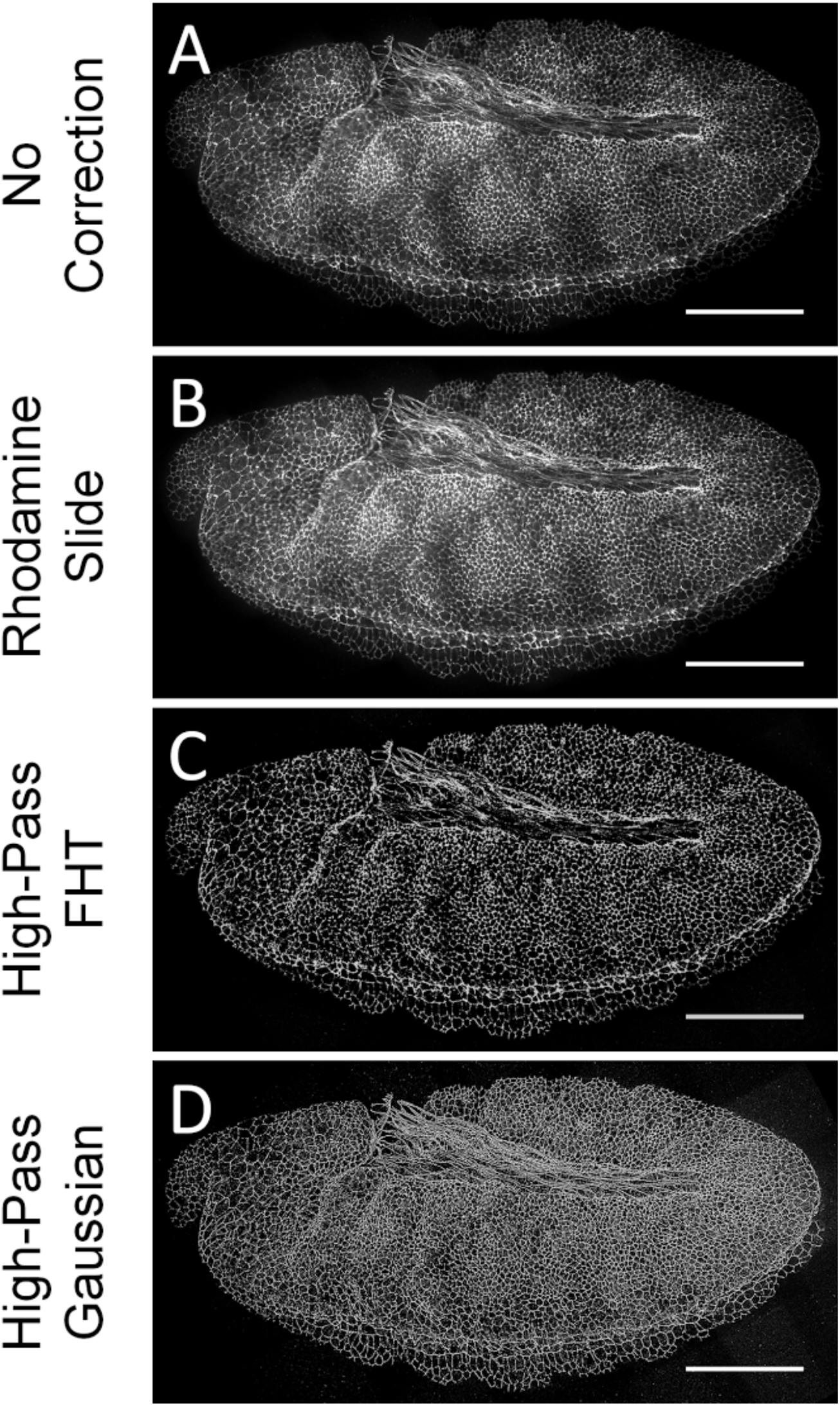
Low-pass filtering with inter-tile normalization results in reduction of inter-tile seams. A-D: stitched 6×5 mosaic of a stage 11 *Drosophila* embryo. A: No correction applied. B: Correction applied via rhodamine correction slide method. C: High-pass FHT filtering and normalization applied. D: High-pass Gaussian filtering and normalization was applied. Scale bar represents 100 μm.

### Confocal microscopy

An Andor spinning disc confocal microscope with a piezo stage at 1.0 μm intervals was used to collect confocal z-stacks. Six by six grids with thirty-three percent overlap were collected for each of the four channels for each embryo. The same settings were used to obtain images of the Rhodamine test slide. Blank images were generated without fluorescent media. MetaMorph^®^ version 7.0.11 was employed for image collection.

### Rhodamine correction slide

A saturated Rhodamine (R6626-25G, Sigma) solution in phosphate-buffered saline was filtered with a syringe filter and placed on a glass slide and sealed with fingernail polish (wet n wild, Wild Shine).

### FFT filtering

Prior to normalization, images were processed to remove vignetting artifacts. A maximum-intensity z-projection of each z-stack was generated using Miji (22), a MATLAB implementation of Fiji (23). For the dpERK antibody stainings, a median filter of radius 3 pixels was employed to remove salt-and-pepper noise caused by the low intensity of the signal relative to the background. The FFT high-pass filter in Fiji was utilized to eliminate vignetting artifacts present in individual tiles. A qualitative parameter sweep revealed that a large object filter between 80 and 150 pixels was sufficient to remove the vignetting artifacts while preserving small image features.

### Gaussian Filtering

As an alternative to FFT, images were processed with a high-pass Gaussian filter,

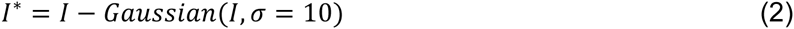

where ***I***\* is the processed image, ***I*** is the raw image, and *σ* is the sigma radius of the Gaussian filter.

### Normalization

To compare or stitch multiple transformed images, objects of similar intensity must first be registered as having the same signal strength. We implement a recursive algorithm to simultaneously normalize adjacent sets of an increasing number of tiles (Figure 1). In each recursion step, we linearly rescale sets of adjacent tiles:

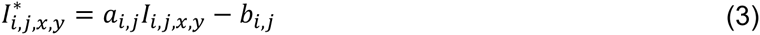

where the parameter matrices *a* and *b* are independent of *x* and *y,* and *i* and *j* represent grid coordinates of tiles within a mosaic. The parameters *a*_*i,j*_ and *b*_*i,j*_ are chosen such that the intensity differences to neighboring tiles are minimized in the overlapping region. At each recursion step we consider up to 2×2 sets of adjacent tiles (Figure 1A). We minimize a cost function using global search algorithm employing the *fmincon* local solver in MATLAB R2015a in order to minimize the intensity differences in the overlap (24). The cost function takes different forms depending on whether the overlap contains two (2×1 or 1×2) or four (2×2) tiles. For the case of the two adjacent sets of tiles the cost function is:

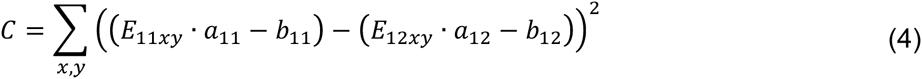

where *E*_11*xy*_ and *E*_12*xy*_ denote the intensity of pixels at the edges of the two overlapping tiles, while *a_ij_* and *b*_*i,j*_ are the optimized parameters for their respective tiles. The resulting parameters are used to modify the tiles using Eq. 2. Importantly, the optimization underlies the constraints that *a* must be greater than one, and *b* must be greater than zero in order to avoid negative intensities, and reduce losses in precision. The cost function for the remaining cases is constructed similarly. To reduce the computational cost of the optimization, intensities in Eq. 4 are binned to one-fourth resolution.

**Figure 1:**
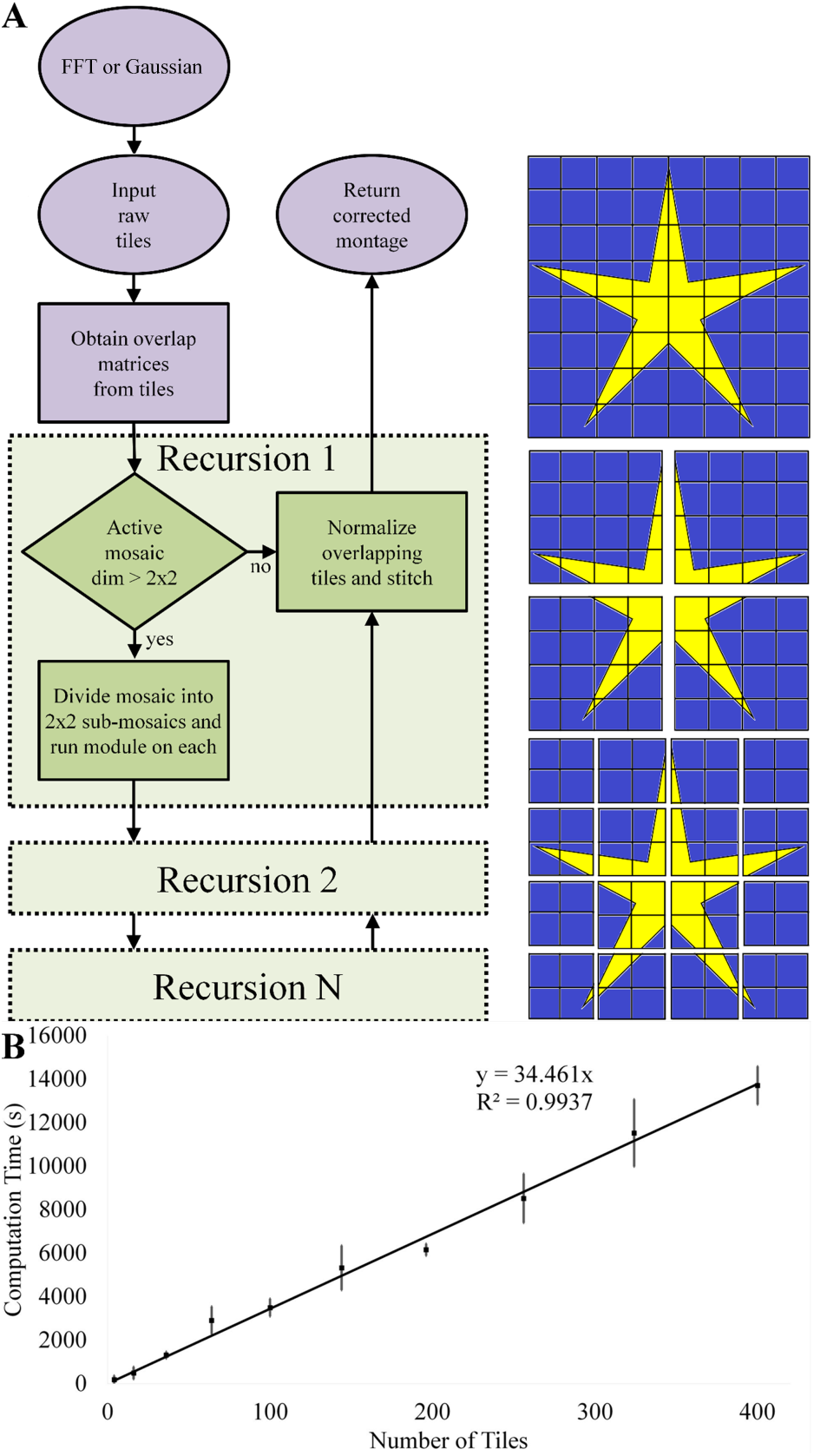
Overview of image processing method. A) The flow diagram shows how a multi-tile mosaic is processed after vignetting correction to normalize tiles and enable subsequent stitching into a mosaic. Mosaics of size 2×2 or smaller are optimized to minimize the difference in the overlaps, as defined by Eq. 3. Mosaics larger than 2×2 are subdivided into sub-mosaics in a 2×2 configuration, and each sub-mosaic is processed in a recursive instance of the script. The normalized sub-mosaics are then normalized through the 2×2 optimization I using Eq. 3. As a result, no optimization is ever I run with more than six independent parameters, and computation time of mosaics of any size) scales nearly linearly with the number of 1 overlaps. B) The correction was run on montages with dimensions MxM, with M ranging ן from 2 to 20 in multiples of two, where 6 (or a total of 36 tiles) represents the dataset tested in this study. The image used contained a linear gradient, divided into tiles with 10% overlap. The tiles were multiplied by a fourth order cosine function to simulate vignetting, and Gaussian noise with a standard deviation of 10% was applied. The resulting data showed a linear correlation between the number of tiles stitched and computation time. N = 5 for each condition, error bars represent standard deviation.

### Algorithm testing

We validated the image processing pipeline on 24 late-stage *Drosophila* embryos expressing CD8::GFP under the *engrailed* enhancer. Engrailed is expressed in the posterior compartments of embryonic segments (25). This marker was chosen to test the method because bands of Engrailed expression are frequently larger than one field of view, allowing the effect of QuickStitch on large features to be studied. IHC was used to visualize cell boundaries (using DE-cadherin) in order to obtain cell-level morphological information and assess the ability of QuickStitch to correct vignetting in cases of small details. Further, measurements of gene regulatory molecules are important for understanding the mechanisms behind cell-signaling and other developmental processes. Doubly phospho-ERK (dpERK), the downstream target of the Epidermal Growth Factor Receptor (EGFR) pathway, is important for many developmental functions including embryonic compartment size homeostasis (26,27). dpERK was assayed, corrected and stitched in order to assess the ability of QuickStitch to correct vignetting of non-homogeneous multi-cellular features that typically is at a lower intensity.

## RESULTS AND DISCUSSION

Our recursive algorithm linearly normalizes adjacent transformed tiles to minimize the intensity difference of overlap pixel intensities (Figure 1A). The recursive nature of the algorithm results in linear scaling of the calculation time with sample size (Figure 1B). High-pass filtering is used by this application to decompose raw images into a summation of component images characterized by differing frequencies. Normalization is needed because the high-pass filtering determines the shape of the vignetting artifact, but not the absolute intensity.

The gold standard Rhodamine correction slide method did not remove vignetting artifacts from *Drosophila* embryo mosaics (Figure 2). QuickStitch was qualitatively more effective at removing vignetting artifacts from *Drosophila* embryo mosaics in three fluorescent channels with very different signals (Figure 3). By removing vignetting artifacts from cell-boundary images (Figure 3C′), watershed segmentation applications such as Seedwater Segmenter (28) and custom segmentation software could quickly be used to segment cell boundaries for the whole embryo with fewer errors in automatic seed selection, as cellular minima occur in a narrower spatial range, and with fewer errors in boundary identification, as cell boundaries will exist in a narrower intensity range. The *en>GFP* signal (Figure 3B), which represents lineage-restricted compartments in the embryo, is an example of a large-scale pattern or structure within the tissue useful for cell classification. Cell-fate classification by thresholding *en* is dependent on a uniform signal, and will result in fewer classification errors with this correction (Figure 3B′). Even features with low intensities such as the spatial pattern of dpERK antibody staining (Figure 3A) were preserved through the normalization process. By correcting for vignetting, the pattern of dpERK activity across multiple tiles is much more apparent to the eye, and matches previous low resolution reports of dpERK expression (20,26). For example, after correction, the characteristic tracheal pit pattern is visible (Figure 2E, 2E′,(26)). This demonstrates the value of vignetting correction for observing phenomenon where the magnitude of the signal is comparable to the magnitude of vignetting artifacts such as high-magnification imaging of the low-intensity gradient of dpERK.

**Figure 3:**
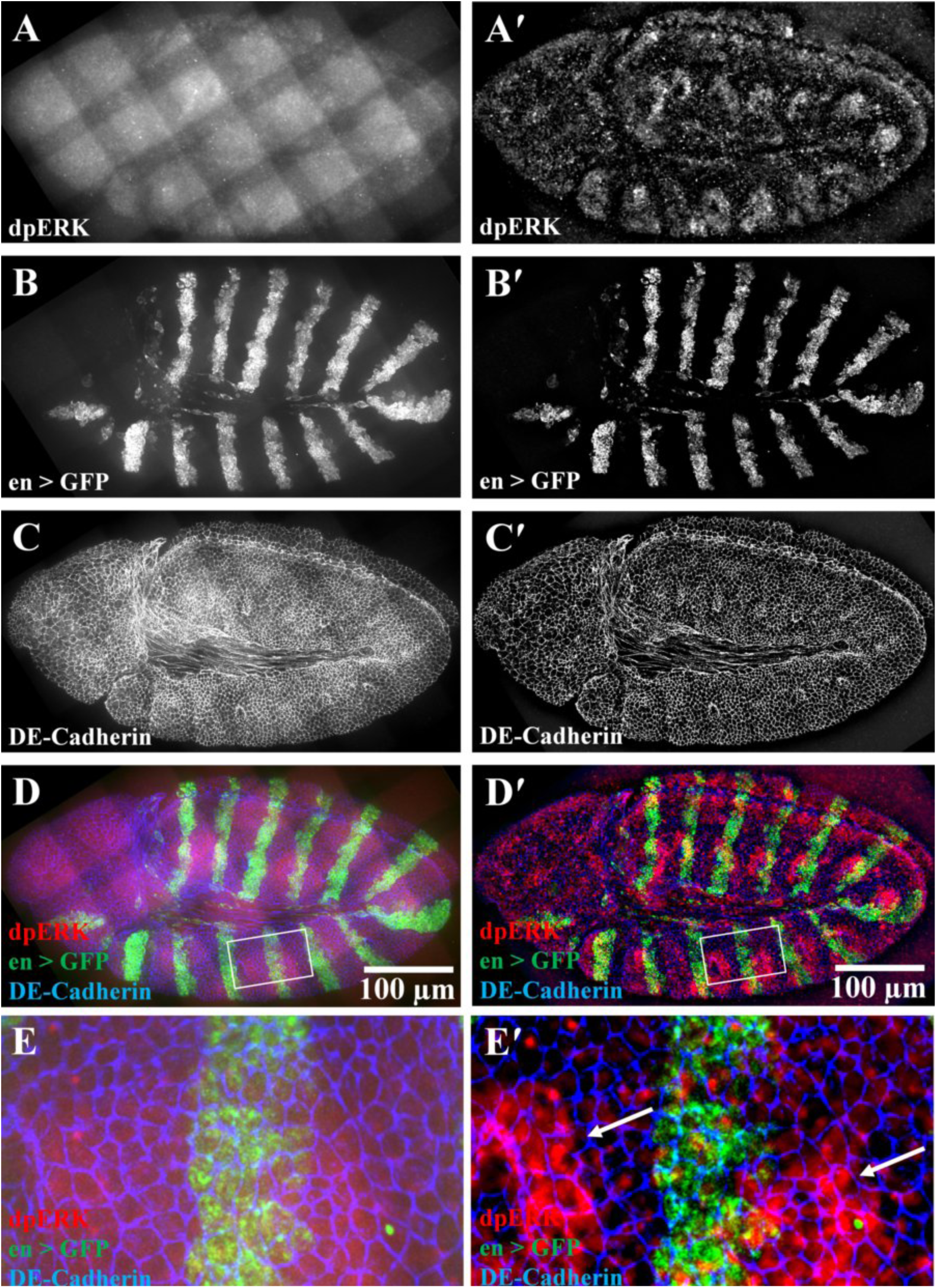
Vignetting correction of a multiplexed immunofluorescent-stained Drosophila embryo. (A-E) Uncorrected and stitched confocal z-projections of a stage-11 *Drosophila* embryo. All three channels show vignetting. (A) For dpERK, which has a low signal, the effect is very strong. (B) Engrailed is localized in the posterior compartment of segments. (C) DE-Cadherin marks the boundaries of cells. (A′-E′) The same images, corrected and normalized using our QuickStitch algorithm. Images have a resolution of 2436×1284 pixels, allowing for high precision cell-level analysis. (E′) Arrows represent the characteristic tracheal pit pattern where dpERK is highly concentrated at this stage of development that is revealed by QuickStitch.

QuickStitch is efficient and preferable over background segmentation algorithms in cases where the tissue makes up the majority of the field of view or where little background information is available such as histological imaging or confluent cell cultures. QuickStitch relies on the assumption that vignetting is a low-frequency phenomenon, and hence can be removed by using a FFT high-pass filter. A basic assumption of this method is that features are smaller than the cutoff frequency. This implementation was developed for two-dimensional z-projections of confocal stacks where z-slices do not have an overlap in the z direction. QuickStitch is open source (Supplemental Code), and is available online as a pre-compiled executable for Macintosh OS X Yosemite and Windows 8.1 operating systems and does not require a MATLAB license to use at: <u>https://notredame.box.com/s/jib2qh1il10jerelhd8irmg0ghqno1bs</u>.

## ACKNOWLEDGEMENTS

We thank Miranda Burnette, Erin Howe, and Qinfeng Wu for comments on earlier versions of the manuscript. We also wish to thank the Bloomington Stock Center for fly lines and Developmental Studies Hybridoma Bank for antibodies. The authors gratefully acknowledge the Notre Dame Integrated Imaging Facility. CN, PB and JZ were funded in part by NSF Award CBET-1403887 and CBET-1553826. PB was funded partially by NIH grant UO1 HL116330. AF and JZ acknowledge funding from a Royal Society International Exchanges Scheme grant (IE130149). JK is funded by the Engineering and Physical Sciences Research Council through a studentship. PB is funded by Walther Cancer Foundation Interdisciplinary Interface Training Project. CN was funded by Notre Dame Advanced Diagnostics & Therapeutics Berry Fellowship.

## AUTHOR CONTRIBUTIONS

PB, CN, JK, AF, and JZ initiated the project and conceived of method. PB collected image data. PB and PE implemented method and performed analysis. PB, PE, and JZ wrote manuscript.

## SI INFORMATION: Supplemental code

~~~
% gridStitchingf.m
% Pipeline for vignetting algorithm % May 8, 2014
% dialogue box with multiple inputs
function gridStitchingf(input, output, outputfilename, M, N, x, y, …
   OverlapPercent, CropPercent, numembs, numberSize, fluor,…
   lowerBounds, upperBounds, x0, MFE, MI, …
   TolCon, TolFun, TolX, timeLimit, ST1, Trial, s)
parpool('local’);
%addpath([filesep ‘Applications’ filesep ‘Fiji.app’ filesep ‘scripts’])
%addpath([’scripts’])
[executionPath,~,~] = fileparts(mfilename(’fullpath’));
javaaddpath ([executionPath filesep ‘mij.jar’]);
tempPath = [executionPath filesep ‘temp’];
mkdir(tempPath);
Miji(false);
% add panel with default java path, should have matlab installation path
% with option to change it
%javaaddpath
’/Users/peberts/Documents/MATLAB/matlab_R2014b_maci64/java/mij.jar’;
%javaaddpath
’/Users/peberts/Documents/MATLAB/matlab_R2014b_maci64/java/ij.jar’;
%MIJ.start;
[pathstr,name,ext] = fileparts(input);
tf = false;
x = length(name);
cc1 = strfind(name, ‘{’);
name_1 = name(1:cc1-1);
cc2 = strfind(name, ‘}’);
name_2 =
name(cc2+1:x);
i_num = cc2-cc1-1;
mkdir([output, filesep, outputfilename])
for k = 1:numembs
%    x = 1;
%       while tf == 0
%          tf = isstrprop(name(x), ‘digit’);
%          x = x + 1;
%       end
%
%    name2 = name(1:x-2);
  channum = max(size(fluor)); for m = !:channum,•
     for i = 1:M
      for j = 1:N
         %for s = s:((M*N))
         name_3 = [name_1, sprintf([’%.’, num2str(i_num), ‘d’], s), ‘_w’, num2str(m), ‘Confocal ‘, num2str(fluor(m)),’_MIP’];
%% Call FFT function
         inc_path = [pathstr, filesep, name_3, ext];         b = [’path=[’, inc_path, ‘]’];
         MIJ.run(’Open…’, b);
         c = [’filter_large=100 filter_small=0 suppress=None
tolerance=5’];
         MIJ.run(’Bandpass Filter…’, c);
         temp = MIJ.getCurrentImage;
         rawTiles(i, j, 1:size(temp,1),1:size(temp,2),m) =
uint16(temp);
         if m == 1
            MIJ.run(’Median…’, ‘radius=3’);
            rawTiles(i,j,:,:,m) = MIJ.getCurrentImage;
         end
         MIJ.run(’Close All’);
         s = s+1;
      end
   end
%% Call Optimization
   r = rawTiles(:,:,:,:,m);
   Global_Stitching_Optimization(r, m, M, N, x, y, OverlapPercent,
CropPercent, …
   input, lowerBounds, upperBounds, x0, MFE, MI, TolCon, TolFun, TolX,
   timeLimit, ST1, Trial, name_1, numembs, s, tempPath, …
   numberSize, channum, fluor, k);
%% Call Stitching
   q = 1;
      b = [’type=[Grid: row-by-row] order=[Right & Down] grid_size_x=’ int2str(M),’ grid_size_y=’ int2str(N), ‘ tile_overlap=’ num2str(OverlapPercent*100),’ first_file_index_i=1 directory=[’ tempPath, ‘] file_names=[’, name_1, ‘Channel - ‘, num2str(fluor(m)), ‘ Tile - {iiii}.tif] output_textfile_name=TileConfiguration.txt fusion_method=[Linear Blending] regression_threshold=0.30 max/avg_displacement_threshold=2.50 absolute_displacement_threshold=3.50 subpixel_accuracy computation_parameters=[Save memory (but be slower)] image_output=[Fuse and display]’];
      MIJ.run(’Grid/Collection stitching’, b);
      temp = MIJ.getCurrentImage;
      stitchedImage(1:size(temp,1),1:size(temp,2),m) = temp;
      MIJ.run(’Close All’);
      Composite(:,:,k,m) = (stitchedImage(:,:,m));
      rmdir(tempPath, ‘s’);
      mkdir(tempPath);
   s = s-(M*N);
  end
 s = s+(M*N);
end
  padding = 150;
  [sizex, sizey, sizez, sizet] = size(Composite);
  Composite2(padding:(sizex+padding-1),padding:(sizey+padding-1),:,:) =
Composite;
  Composite2(padding*2+sizex,padding*2+sizey,sizez,sizet) = 0;
  Composite = Composite2;
  clear Composite2;
  for m = 1:channum
   a = double(max(max(Composite(:,:,:,m))));
   c = double(Composite(:,:,:,m));
   b = (c./a)*65536;
   s = [output, filesep, outputfilename, ‘ emb ‘, num2str(fluor(m)),
’.tif’];
   imwrite(uint16(b), [s]);
  end
delete(gcp)
end
~~~

### Primary recursive algorithm

~~~
% Hand this function an M * N * x * y * 4 set of edge intensities (E) and
% the function will make several decisions: if the matrix is sufficiently % small (2×1, 1×2, 1×1) it will directly call the optimization wrapper and
% return a and b parameters in the form E* = aE - b that minimizes the
% difference between corresponding E matricies. If the matrix is larger, it % recursively calls itself for each half of the matrix, obtains a and b
% parameters, obtains E1* and E2* from the a, b and E values and returns
% new parameters. This algorithm should not modify E1, E2, x and y global % variables, but instead call the wrapper to do so.
% Unlike the optimization wrapper, this function returns an M x N matrix of
% parameters for a and b. This function does not handle binning of edge
% matricies, that is done in the main function.
% try adding a label to the a and b values to pass throughout the
% normalization
function [ a, b ] = a_b_recursive( E )
%% Obtain the size parameters from input matrix
[M, N, ~, y, ~] = size(E); %Determine tile layout
xlength = round((M + 0.5) / 2); %Number of tiles if split in x direction
ylength = round((N + 0.5) / 2); %Number of tiles if split in y direction
if xlength > 2
   xlength = xlength - mod(xlength, 2);
end
if ylength > 2
   ylength = ylength - mod(ylength, 2);
end
%% Determine which algorithm to run depending on M x N dimensions of input matrix
if M == 2 && N == 2
   qq = E(1,1,:,:,2);
   ww = E(2,1,:,:,3);
   ee = E(2,2,:,:,4);
   rr = E(1,2,:,:,1);
   tt = E(2,1,:,:,4);
   yy = E(2,2,:,:,1);
   uu = E(1,2,:,:,2);
   ii = E(1,1,:,:,3);
   [ a, b ] = optimize_a_b_2×2(qq,ww,ee,rr,tt,yy,uu,ii);
elseif M == 2 && N == 1 % Run optimization for 2×1
   [ a, b ] = optimize_a_b(E(1,1,:,:,2), E(2,1,:,:,4));
   a = a’;
   b = b’;
elseif M == 1 && N == 2 % Run optimization for 1×2
[ a, b ] = optimize_a_b(E(1,1,:,:,3), E(1,2,:,:,1));
elseif M == 1 && N == 1 % Optimization is finished for this tile
   a = 1;
   b = 0;
else
   [temp_a11, temp_b11] = a_b_recursive(E(1:xlength,1:ylength,:,:,:)); %
Obtain parameters for left side tiles
   [temp_a21, temp_b21] = a_b_recursive(E((xlength+1):M,1:ylength,:,:,:)); %
Obtain parameters for right side tiles
   [temp_a12, temp_b12] = a_b_recursive(E(1:xlength,(ylength+1):N,:,:,:)); %
Obtain parameters for left side tiles
   [temp_a22, temp_b22] =
a_b_recursive(E((xlength+1):M,(ylength+1):N,:,:,:)); % Obtain parameters for right side tiles
   % Overlap 1 (vertical top)
   for j = 1:ylength
      left_temp(:,:) = E(xlength,j,:,:,2) * temp_a11(xlength,j) - temp_b11(xlength,j);
      right_temp(:,:) = E(xlength+1,j,:,:,4) *
temp_a21(1,j) - temp_b21(1,j);
      qq (:,((j-1)*y+1):(j*y)) = left_temp;
      tt (:,((j-1)*y+1):(j*y)) = right_temp;
   end
   % Overlap 2 (horizontal right)
   for i = 1:(M-xlength)
      top_temp(:,:) = E(i+xlength,ylength,:,:,3) * temp_a21(i,ylength) - temp_b21(i,ylength);
      bottom_temp(:,:) = E(i+xlength,ylength+1,:,:,1) * temp_a22(i,1) - temp_b22(i,1);
      ww (:,((i-1)*y+1):(i*y)) = top_temp;
      yy (:,((i-1)*y+1):(i*y)) = bottom_temp;
   end
   % Overlap 3 (vertical bottom)
   for j = 1:(N-ylength)
   left_temp(:,:) = E(xlength,j+ylength,:,:,2) * temp_a12(xlength,j) - temp_b12(xlength,j);
   right_temp(:,:) = E(xlength+1,j+ylength,:,:,4) * temp_a22(1,j) -
temp_b22(1,j);
      uu (:,((j-1)*y+1):(j*y)) = left_temp;
      ee (:,((j-1)*y+1):(j *y)) = right_temp;
   end
   % Overlap 4 (horizontal left)
   for i = 1:xlength
      top_temp(:,:) = E(i,ylength,:,:,3) * temp_a11(i,ylength) - temp_b11(i,ylength);
      bottom_temp(:,:) = E(i,ylength+1,:,:,1) * temp_a12(i,1) -
temp_b12(i,1);
      rr (:,((i-1)*y+1):(i*y)) = bottom_temp;
      ii (:,((i-1)*y+1):(i*y)) = top_temp;
   end
   [super_a, super_b] = optimize_a_b_2×2(qq,ww,ee,rr,tt,yy,uu,ii);
   a(1:xlength,1:ylength) = temp_a11 * super_a(1,1);
   a((xlength+1):M,1:ylength) = temp_a21 * super_a(2,1);
   a(1:xlength,(ylength+1):N) = temp_a12 * super_a(1,2);
   a((xlength+1):M,(ylength+1):N) = temp_a22 * super_a(2,2);
   b(1:xlength,1:ylength) = temp_b11 * super_a(1,1) + super_b(1,1);
   b((xlength+1):M,1:ylength) = temp_b21 * super_a(2,1) + super_b(2,1);
   b(1:xlength,(ylength+1):N) = temp_b12 * super_a(1,2) + super_b(1,2);
   b((xlength+1):M,(ylength+1):N) = temp_b22 * super_a(2,2) + super_b(2,2);
end
~~~

### Optimization function for 2×1 overlap

~~~
% This function returns the fitness of the solution for a combination of
% parameters. It should not modify any global variables, but reads E from
% the optimization wrapper and dimensions from the recursive algorithm
% call. SA is the solution matrix in the form: [a11 a12 b11 b12].
% SA: [a1/a2 b1-b2]
function [ SSR ] = optimization_overlaps( SA )
   global E1 E2;
   SSR = sum(sum((E1 * SA(1) - E2 * SA(2) - SA(3) + SA(4)).^2));
end
~~~

### Wrapper for 2×1 overlap

~~~
% The purpose of this function is to serve as a wrapper for the
% optimization problem of two overlapping regions (E1 and E2). The function
% takes in two regions and returns the a and b values which will result in
% the most seamless fit of the two images. No other function should modify
% E1 and E2. This function returns a and b as arrays of length 2.
function [ a, b ] = optimize_a_b(matrix1, matrix2)
   global E1 E2 output;
   global lb ub x0 MFE MI TolCon TolFun TolX timeLimit ST1 Trial
   [row, ~] = size(output);
   E1 = matrix1;
   E2 = matrix2;
   problem.objective = @optimization_overlaps;
   problem.nvars = 4;
   problem.lb = [ones(1,2)*lb(1) ones(1,2)*lb(2)];
   problem.ub = [ones(1,2)*ub(1) ones(1,2)*ub(2)];
   problem.solver = ‘fmincon’;
   problem.x0 = [ones(1,2)*x0(1) ones(1,2)*x0(2)];
   problem.options = optimoptions(’fmincon’,’Display’,’none’);
   problem.options.MaxFunEvals = MFE;
   problem.options.MaxIter = MI;
   problem.TolCon = TolCon;
   problem.TolFun = [ones(1,2)*TolFun(1) ones(1,2)*TolFun(2)];
   problem.TolX = TolX;
   gs =
GlobalSearch(’MaxTime’,timeLimit,’NumStageOnePoints’,ST1,’NumTrialPoints’,Tri al, ‘Display’, ‘off’);
   [solution,fval,flag,~,solutions] = run(gs, problem);
   output{row+1,1} = fval;
   output{row+1,2} = solutions;
   disp([’2×1 GS finished with flag: ‘ num2str(flag) ‘ GS run: ‘
num2str(row+1)]);
   a = reshape(solution(1:2),[1, 2]);
   b = reshape(solution(3:4),[1, 2]);
   %if solution(1) > 1
   %    a = [solution(1), 1];
   %else
   %    a = [1 1/solution(1)];
   %end
   %if solution(2) >
   %    b = [solution(2), 0];
   %else
   %    b = [0, -solution(2)];
   %end
end
~~~

### Optimization function for 2×2 overlap

~~~
% This function returns the fitness of the solution for a combination of
% parameters. It should not modify any global variables, but reads E from
% the optimization wrapper and dimensions from the recursive algorithm
% call. SA is the solution matrix in the form: [a11 a12 b11 b12].
% SA: a, then b in form [11 21 12 22]
function [ SSR ] = optimization_overlaps_2×2( SA )
   global E11 E12 E13 E14 E21 E22 E23 E24;
   SSR = sum(sum((E11 * SA(1) - E21 * SA(2) - SA(5) + SA(6)).^2));
   SSR = SSR + sum(sum((E12 * SA(2) - E22 * SA(4) - SA(6) + SA(8)).^2));
   SSR = SSR + sum(sum((E13 * SA(4) - E23 * SA(3) - SA(8) + SA(7)).^2));
   SSR = SSR + sum(sum((E14 * SA(3) - E24 * SA(1) - SA(7) + SA(5)).^2));
end
~~~

### Wrapper for 2×2 overlap

~~~
% The purpose of this function is to serve as a wrapper for the
% optimization problem of two overlapping regions (E1 and E2). The function
% takes in two regions and returns the a and b values which will result in
% the most seamless fit of the two images. No other function should modify
% E1 and E2. This function returns a and b as arrays of length 2.
function [ a, b ] = optimize_a_b_2×2(q, w, e, r, t, y, u, i)
   global E11 E12 E13 E14 E21 E22 E23 E24 output;
   global lb ub x0 MFE MI TolCon TolFun TolX timeLimit ST1 Trial
   E11 = q;
   E12 = w;
   E13 = e;
   E14 = r;
   E21 = t;
   E22 = y;
   E23 = u;
   E24 = i;
   [row, ~] = size(output);
   problem.objective = @optimization_overlaps_2×2;
   problem.nvars = 8;
   problem.lb = [ones(1,4)*lb(1) ones(1,4)*lb(2)];
   problem.ub = [ones(1,4)*ub(1) ones(1,4)*ub(2)];
   problem.solver = ‘fmincon’;
   problem.x0 = [ones(1,4)*x0(1) ones(1,4)*x0(2)];
   problem.options = optimoptions(’fmincon’,’Display’,’none’);
   problem.options.MaxFunEvals = MFE;
   problem.options.MaxIter = MI;
   problem.TolCon = TolCon;
   problem.TolFun = [ones(1,4)*TolFun(1) ones(1,4)*TolFun(2)];
   problem.TolX = TolX;
   gs =
GlobalSearch(’MaxTime’,timeLimit,’NumStageOnePoints’,ST1,’NumTrialPoints’,Tri al,’Display’,’off’);
   [solution,fval,flag,~,solutions] = run(gs, problem);
   output{row+1,1} = fval;
   output{row+1,2} = solutions;
   disp([’2×2 GS finished with flag: ‘ num2str(flag) ‘ GS run: ‘
num2str(row+1)]);
   a = reshape(solution(1:4),[2 2]);
   b = reshape(solution(5:8),[2 2]);
end
~~~

